# Deep Learning on Multimodal Chemical and Whole Slide Imaging Data for Predicting Prostate Cancer Directly from Tissue Images

**DOI:** 10.1101/2022.05.11.491570

**Authors:** Md Inzamam Ul Haque, Debangshu Mukherjee, Sylwia A. Stopka, Nathalie Y.R. Agar, Jacob Hinkle, Olga S. Ovchinnikova

## Abstract

Prostate cancer is one of the most common cancers globally and is the second most common cancer in the male population in the US. Here we develop a study based on correlating the H&E-stained biopsy data with MALDI mass-spectrometric imaging of the corresponding tissue to determine the cancerous regions and their unique chemical signatures, and variation of the predicted regions with original pathological annotations. We spatially register features obtained through deep learning from high-resolution optical micrographs of whole slide H&E stained data with MSI data to correlate the chemical signature with the tissue anatomy of the data, and then use the learned correlation to predict prostate cancer from observed H&E images using trained co-registered MSI data. We found that this system is more robust than predicting from a single imaging modality and can predict cancerous regions with *∼*80% accuracy. Two chemical biomarkers were also found to be predicting the ground truth cancerous regions. This will improve on generating patient treatment trajectories by more accurately predicting prostate cancer directly from H&E-stained biopsy images.

## 1 Introduction

Prostate cancer (PC) is the second most common cause of cancer as well as the second leading cause of cancer death among men. Approximately 1 in 8 men will be diagnosed with PC during their lifetime [1]. Therefore, a significant amount of effort has been focused on developing novel treatment options [2], early detection [3] as well as predictive models for risk prediction [4]. In particular, work has focused on forecasting metastatic PC (mPC) [5] as it is most likely to lead to patient death. A lot of work have focused on developing artificial intelligence (AI) and machine learning (ML) approaches to automatically grade hematoxylin and eosin (H&E) pathology slides to address these problems [6], [7], [8], [9], and [10]. The first statistical model to predict a gene mutation in cancer directly from the patient’s digitized H&E-stained whole microscopy slide was presented in [11].

Mass spectrometry imaging (MSI) offers a path to improve detection and labeling of pathology slides by introducing a chemical imaging modality that is able to identify mPC based on chemical biomarkers with much higher confidence [12], [13]. There have been multiple studies regarding the identification/diagnosis of prostate cancer using mass spectrometry (MS). For example, Andersen et al [14] showed that specific tissue compartments within prostate cancer samples have distinct metabolic profiles and using matrix assisted laser desorption ionization-time of flight (MALDI-TOF) MSI data they identified several differential metabolites and lipids that have potential to be developed further as diagnostic and prognostic biomarkers for prostate cancer. Various MALDI-based MS techniques including imaging, profiling and proteomics in-depth analysis where MALDI MS follows fractionation and separation methods such as gel electrophoresis, have been used to identify prostate cancer biomarkers [15]. A more recent study [16] found nine key biomarkers when using MSI on intact human prostate tissue specimens that determined metabolites which could either differentiate between benign and malignant prostate tissue or indicate prostate cancer aggressiveness. Therefore, work on the integration of multimodal bioinformation data, specifically MSI and H&E data offers the potential to improve on identification and accurate labeling of PC. In 2015, [17] reported a data fusion framework for MSI and H&E stain microscopy enabling prediction of a molecular distribution both at high spatial resolution and with high chemical specificity. [18] compared two pansharpening methods, Intensity–Hue–Saturation and Laplacian Pyramid, and demonstrated the latter was more robust for image fusion between MSI and electron microscopy. However, these fusion based approaches are limited by the fundamental difference between physical mechanisms of image generation for MSI and techniques used for data up-sampling and are prone to reconstruction errors and are unable to reconstruct full spectral predictions. To circumvent these problems our group has demonstrated that a physically constrained model between two MSI imaging techniques is able to accurately reconstruct and predict both high spatial resolution images as well as high spectral resolution mass spectra [19].

In this work we demonstrate a machine learning approach that utilizes whole slide H&E pathology labeled data and MSI data from a 9.4T MALDI Fourier-transform ion cyclotron resonance (FTICR) MS for predicting PC directly from H&E tissue images. We show, as is reported in previous literature [12] that MSI is extremely useful for predicting cancerous regions, and we leverage this to develop a pipeline for predicting cancerous regions using both WSI-to-MSI and MSI-to-PC stages. Since MSI is expensive and not widely available, our approach uses paired MSI and WSI data which does not require manual labels to train the WSI-to-MSI model. The resulting model provides not only a binary cancer/non-cancer segmentation but a spectral estimate at each point of an H&E image, resulting in a highly interpretable prediction. This work lays the groundwork for developing more accurate MSI prediction with larger paired MSI/WSI datasets, and minimizing the amount of manually labelled images required for PC prediction by leveraging the richness of intermediate WSI supervision.

## 2 Materials and Methods

A diagram demonstrating the steps of the proposed approach is shown in Figure 1. It should be noted that there are three stages: only with MSI data, only with H&E data, and multimodal study. We perform model prediction with these three different scenarios and compare them in the results section.

**Figure 1:**
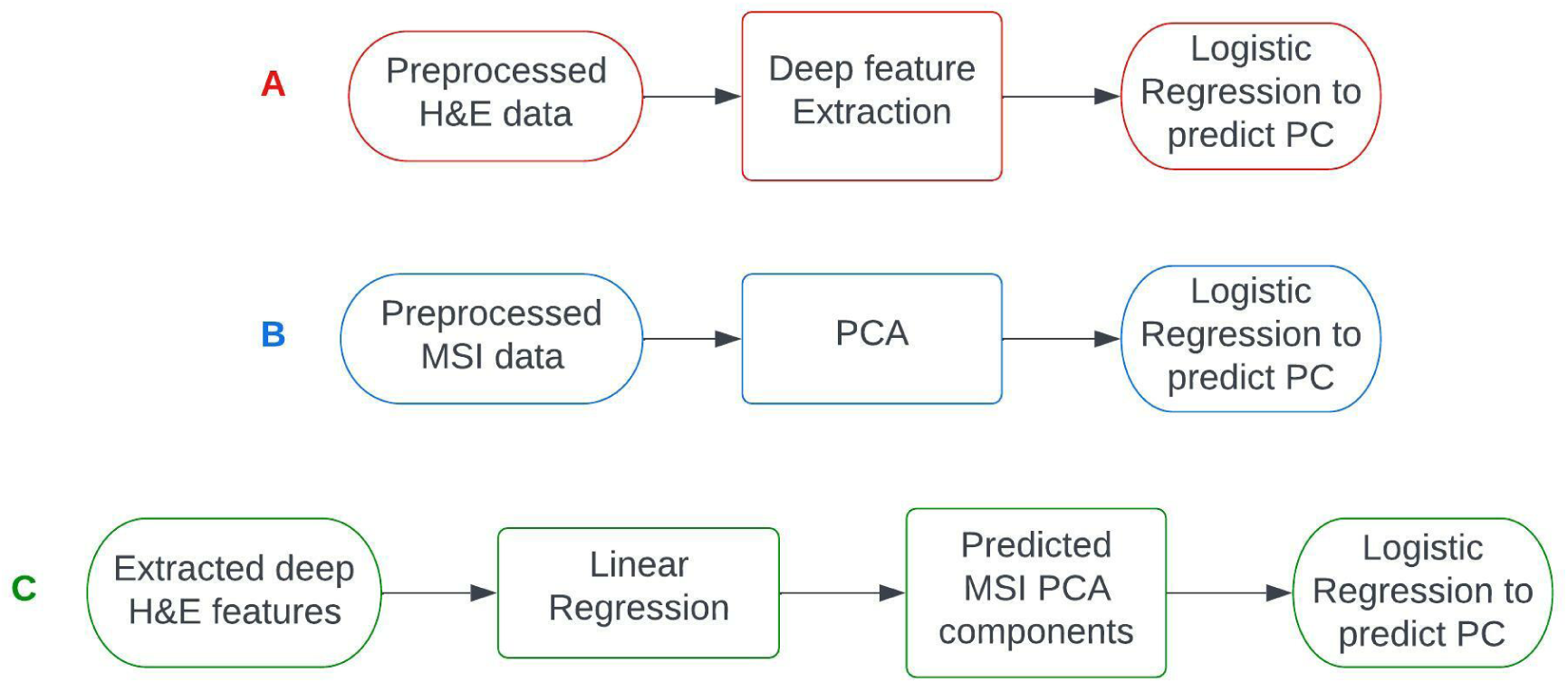
Diagram showing steps of the proposed approach. PC prediction is done with only H&E data (**A**), only MSI data (**B**), and combining both modalities (**C**). Table 1 summarizes the result of this three different approach.

### 2.1 Data

For this study, previously published human prostate RAW files were provided [12] by the authors. Briefly, the mass spectrometry data consisted of human prostate tissue specimens that were cryosectioned and imaged at a pixel size of 120 *µ*m using a 9.4 Tesla SolariX XR FT ICR MS (Bruker Daltonics, Billerica, MA). Corresponding high-resolution annotated H&E images of the same tissue were provided after MALDI matrix removal. Further details regarding data collection could be found in the original publication [12].

In total, we received five tissue samples of MSI data and corresponding annotated high resolution H&E whole slide images. With the five tissue images, a five-fold cross-validation is used during training for all the logistic regression models summarized in Table 1 that are explained later in this section. Hence, the held-out image is part of the five-fold cross-validation. For each fold, total pixels of the four preprocessed tissue images, excluding the heldout image for testing, are shuffled and divided in 80%-20% for training and validation respectively.

**Table 1:**
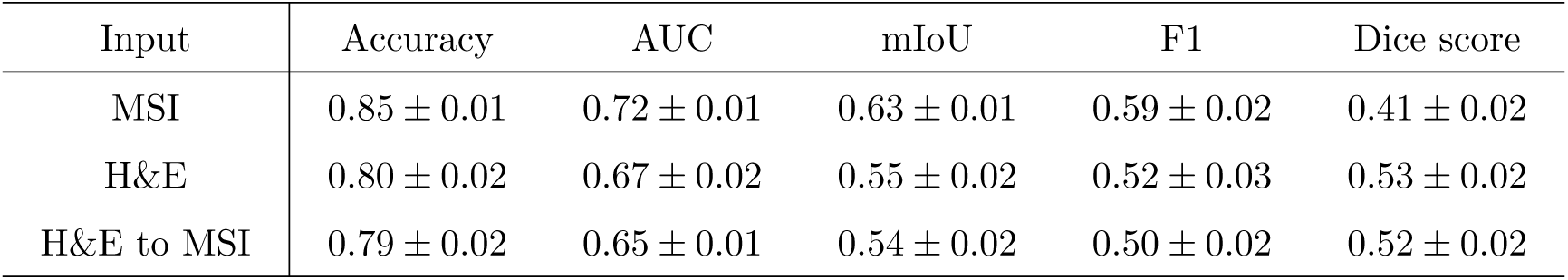
Logistic regression test metrics as mean *±* std using five-fold cross-validation for label prediction from MSI, H&E, and MSI as first predicted from H&E.

### 2.2 MSI data processing

The MSI data was originally acquired in the imzML format - a common data format for MS imaging. To facilitate experimentation, we convert this data to HDF5 format. Since the imzML files are in the range of couple hundred gigabytes, HDF5 format is a suitable choice for fast I/O processing and storage. Each imzML file consists of a large number of mass spectra from which we can produce ion image of the whole slide as well as see spectrum for any spatial coordinate. After inspecting spectrum for several coordinates, we found that there is a slight difference in the *m/z* values between the individual spectrum. To have a common *m/z* axis for all the coordinates in an image, we first interpolate the intensity values for *m/z* values ranging from 100 to 1,000 with a step size of 0.001, and then convert it to HDF5 file format.

### 2.3 Binary mask extraction

Pathologist annotations were provided in the form of contours overlaid low-resolution H&E images. We extract binary masks from the contours for both cancer and non-cancer regions, as well as non-tissue background. These masks are used to register both the high resolution whole slide H&E images and MSI images. Affine transformation followed by phase correlation have been used to co-register H&E and MSI images using the extracted binary masks. After extracting the masks it is possible to see differences in mass spectra between cancer and non-cancer regions.

### 2.4 PCA of MSI data

The interpolated MSI files were too large to conveniently process since they contained 900000 spectral channels at thousands of points. Therefore, we used principal component analysis (PCA) to reduce the dimensionality of each spectrum to 200. These 200 modes of PCA explained 98.5% of the variability in the data with the first three components combined explaining 67% variability (see Figure 2G). Although some modes of PCA captured the cancerous vs. non-cancerous regions better than others, no single PCA mode could clearly segment important regions. Figure 2 shows the first five PCA modes for one of the MSI images. The PCA training was done jointly using the MSI images and tested on a holdout image. Incremental PCA from the scikit-learn library has been used to fit the training in memory and for faster convergence.

**Figure 2:**
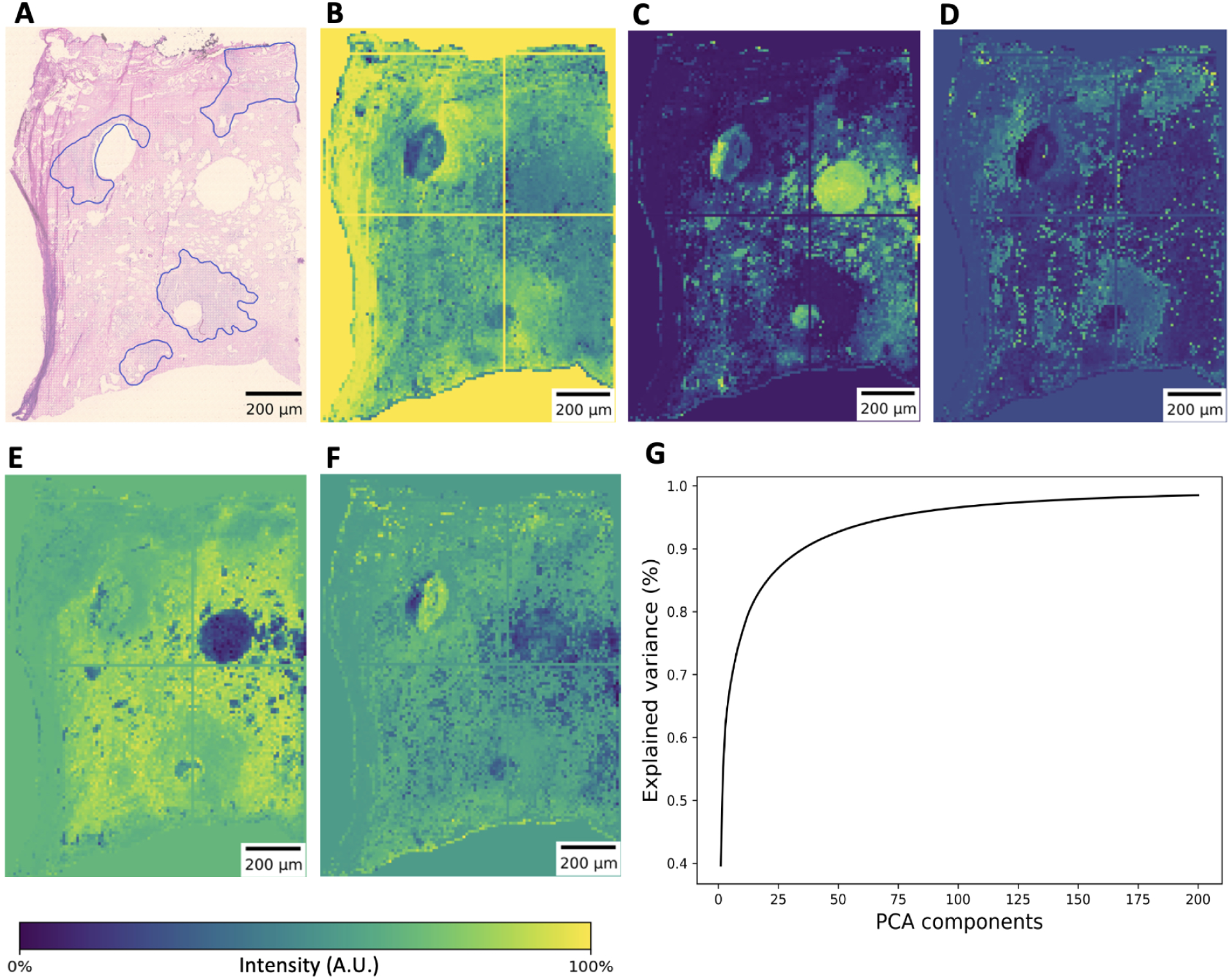
Dimension reduction of MSI using PCA. **(A)** is original cancer annotations by pathologists for a tissue specimen, **(B)** - **(F)** shows first five PCA components respectively for the tissue specimen. A color bar is given showing intensity values in arbitrary units for the five components. Although the components show hints of cancerous regions, no single PCA component can clearly segment cancerous regions. **(G)** shows the cumulative explained variance by the 200 PCA components.

### 2.5 Logistic Regression of MSI

We trained a logistic regression model using the 200 PCA modes of all five MSI images to predict cancerous regions. Five-fold cross validation was used during training. The extracted binary masks were used as the labels for logistic regression. Scikit-learn’s SGD classifier class has been used to perform this regression task. This estimator implements regularized linear models with stochastic gradient descent (SGD) learning. We used the ‘log’ loss that gives logistic regression, a probabilistic classifier. A regularization parameter of 0.01 has been used with an adaptive learning rate starting from 0.01. Since we had a class imbalance in labels, we used a ‘balanced’ class weight fit preset which uses the values of labels to automatically adjust weights inversely proportional to class frequencies in the input data. After training, the model was tested on a holdout image to check performance.

### 2.6 H&E data processing

The high-resolution whole slide H&E tissue images (WSI) were provided in Carl Zeiss CZI data format. Again, for faster I/O processing and storage, we converted this data to HDF5 format. Since these are high-resolution images, we upscaled the binary masks, combine non-cancer and cancer masks to get the foreground and then co-register the H&E images. We also normalized each image by their mean and standard deviation to have zero mean and a unit standard deviation.

### 2.7 MSI prediction from H&E data

First, we extracted features using deep learning from the normalized high resolution WSIs. Inspired from the results of [20], we used a Resnet-50 model for extracting the features from WSIs. Since these are large high-resolution images, we divide each image in 512×512 patches before feeding to the feature extractor. We have also used halos of 256 pixels around the patches to eradicate grid artifacts. This feature extraction step increases the number of channels from 3 to 1024 since we are extracting from the 3rd layer of the Resnet-50 backbone. Also, we reduced the spatial dimension by 16 times using average pooling. This reduction was necessary to get closer to the spatial dimension of the MSI images. Second, to match exactly with the spatial dimension of MSI PCA modes, for each slice, we regridded the extracted features. Considering the extracted features as the source dataset and the MSI PCA modes as the target dataset, we used two strategies: downscaling and upscaling. Downscaling is used when the target shape is is smaller than the source shape. If the source shape is an integer multiple of the target shape, we perform average pooling. Otherwise, we perform Gaussian blurring followed by subsampling. Upscaling is performed when the target shape is larger than the source shape, in which case we use linear interpolation. Having the spatial dimension of both datasets the same, we perform linear regression on the extracted features and compare with the MSI PCA modes. A 5-fold cross validation has been used for training. Adam optimizer with a learning rate of 0.1 has been used, and the convergence took 20000 iterations. We saved the predicted MSI modes and train the same logistic regression model that was used with the H&E data to predict cancerous regions. In other words, we first predict MSI modes from the downscaled H&E features and then train a logistic regression model as discussed in previous subsection with the predicted MSI data to predict cancer labels.

### 2.8 Logistic Regression of H&E data

We trained a logistic regression model using the regridded deep H&E features to predict cancerous regions and to compare performance with the results achieved with MSI data. Again, five-fold cross validation was used during training. The same extracted binary masks that were used for the MSI data training were used as the labels for logistic regression. Scikit-learn’s SGD classifier that implements regularized linear models with stochastic gradient descent (SGD) learning was used for the training. Same as before, we have used the ‘log’ loss that gives logistic regression, a probabilistic classifier. A regularization parameter of 0.001 has been used with an adaptive learning rate starting from 0.001. Similar to the MSI logistic regression training, we used a ‘balanced’ class weight fit preset which uses the values of labels to automatically adjust weights inversely proportional to class frequencies in the input data, essentially removing class imbalance of the training data. After training, the model was tested on a holdout image to see performance.

### 2.9 MSI peak prediction

Previously, [13] identified 32 distinguishable *m/z* signals that occurred in 5–90% of the spectra of in their 729 analyzable tissues samples. They found a total of 15 of these signals appeared to be associated with epithelial structures, based on the comparison with the H&E stain of the slide - *m/z* 605, 616, 644, 678, 700, 899, 976, 1,014,1,044, 1,199, 1,275, 1,502, 3,071, 3,086 and 3,577. We first increased our range of *m/z* values from 1,000 to 1,500 to test most of these spectral features, since our original interpolation accounted for 100-1,000 *m/z* values. Since our H&E to MSI predictions correspond to the MSI PCA modes, we convert the predictions to the original MSI *m/z* spectral dimension, increasing 200 features to 1,500,000 features. Then we choose the above mentioned features in the range from 100 to 1,500 *m/z*. Also, [12] listed *m/z* values that were searched against the Lipid Maps database. We observe ground truth MSI and their prediction from H&E for these *m/z* values. We looked for chemical biomarkers with the ground truth MSI that best captured the cancerous regions of the tissue specimens and then observed the predicted MSI.

## 3 Results

We have trained logistic regression models to predict cancerous regions from MSI PCA modes, regridded H&E data, and converted MSI data from H&E data as discussed in the methods sections. Quantitative results including several metrics are summarized in Table 1. It is a pixel-based result, and there is a class imbalance present. From the total five tissue samples, on average, 14.00 % pixels belong to cancerous regions, 54.78 % pixels belong to non-cancerous regions, and 31.22 % is the background of the images. These results were achieved with a five-fold cross-validation. Here, AUC refers to area under the receiver operating characteristic (ROC) curve, mIoU is the mean intersection over union, also known as the Jaccard similarity coefficient score. The mIoU measures the size of the intersection divided by the size of the union of the true labels and the predicted labels above a given confidence threshold. F1 score can be interpreted as a harmonic mean of the precision and recall of labels above a given threshold. Dice score computes the Dice dissimilarity between predicted and true labels.

From Table 1 we can see that, a better prediction is achieved with MSI data. As previously reported in [12], this result actually shows that PC can be better predicted using chemical information obtained from MSI data. The logistic regression performs similarly when we get label prediction directly from downscaled H&E features and MSI modes predicted from H&E features. Combining H&E and MSI data, we get 79% accuracy with logistic regression. A qualitative result is also shown in Figure 3 which corresponds to the quantitative results we achieved. Predictions for three different tissue specimens are shown in Figure 3. If we look closely in Figure 3B and Figure 3F, some secondary regions are revealed with the MSI to label prediction. These regions can be random noise or cancerous regions missing in the original annotation. When we look at the prediction for the second tissue specimen directly from H&E in Figure 3G, the secondary region seen with MSI prediction is revealed more. Interestingly, this region is revealed even more when we predict from predicted MSI components as seen in Figure 3H. For, other two tissue specimens in Figure 3, cancerous regions are revealed well for all three predictions. Label prediction directly from MSI is best for the third tissue specimen as seen in Figure 3J.

**Figure 3:**
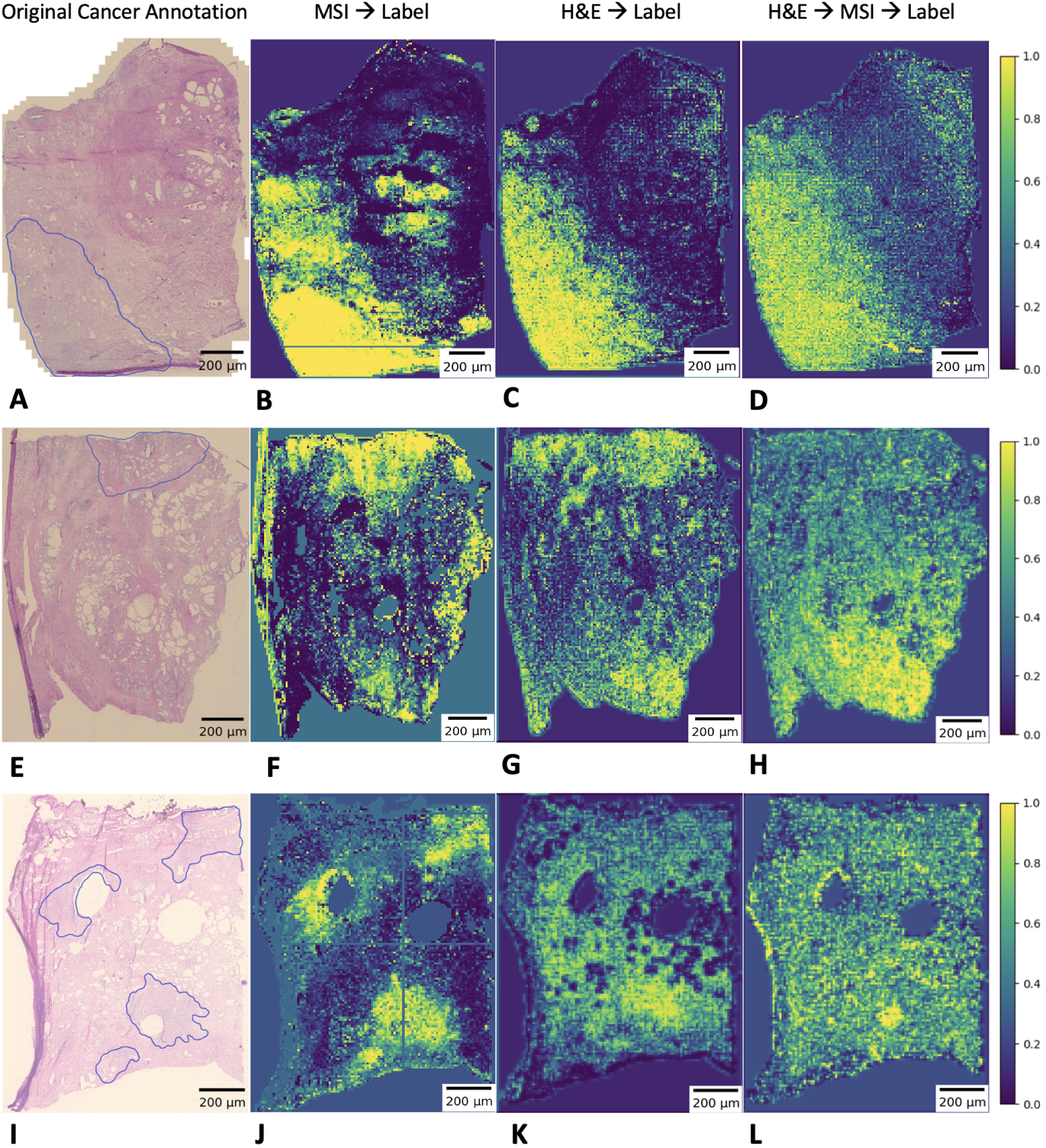
**(A), (E)**, and **(I)** are original cancer annotations for three different tissue specimens. **(B), (F)**, and **(J)** show predicted cancer regions from MSI PCA modes for the three tissue specimens. **(C), (G)**, and **(K)** show predicted cancer labels from regridded H&E features for the three tissue specimens. **(D), (H)**, and **(L)** show prediction of cancer labels from MSI predictions which is predicted form H&E features. All the predictions in this figure are achieved using logistic regression. Row one shows good agreement between MSI **(B)**, and H&E **(C)**, with a possible false positive region that is not labelled in the H&E **(C)** or H&E to MSI **(D)** predicted components. In row two, MSI **(F)**, and H&E **(G)** performs well, but H&E to MSI **(H)** underperforms. In row three, MSI **(J)** significantly outperforms H&E **(K)**, and H&E to MSI **(L)** partially recovers some of that performance.

Figure 4 shows the result of H&E to MSI prediction for three different tissue specimens. As discussed in the methods section, we used a linear regression model with regridded H&E features as the input and MSI PCA modes as the labels. Out of 200 PCA components, prediction for the component having the highest *R*^2^ is shown. We achieved an overall *R*^2^ score of 0.23. In this case, it is expected to not have very high accuracy and *R*^2^ score since we are predicting a different modality of imaging data. Nevertheless, we can see similar patterns in the predicted MSI images corresponding to the actual MSI images. Prediction for tissue specimen 2 as shown in Figure 4B is slightly better compared to the other two tissue specimens. A single pixel is chosen from the cancerous region for each tissue specimen and the mass spectrum is plotted shown in Figure 4C, Figure 4F, and Figure 4I. It is evident that there is an intensity mismatch between the actual and predicted spectrum, but the prediction is clearly able to capture most of the *m/z* peaks in the actual spectrum. Since we interpolated the MSI data in the range of 100 to 1,000 *m/z* values, the corresponding mass spectra are shown in the same range. Also, because of using PCA modes as the labels in this linear regression problem, we ended up having some negative values in the prediction which are clipped to zero in the spectrum plots in this figure.

**Figure 4:**
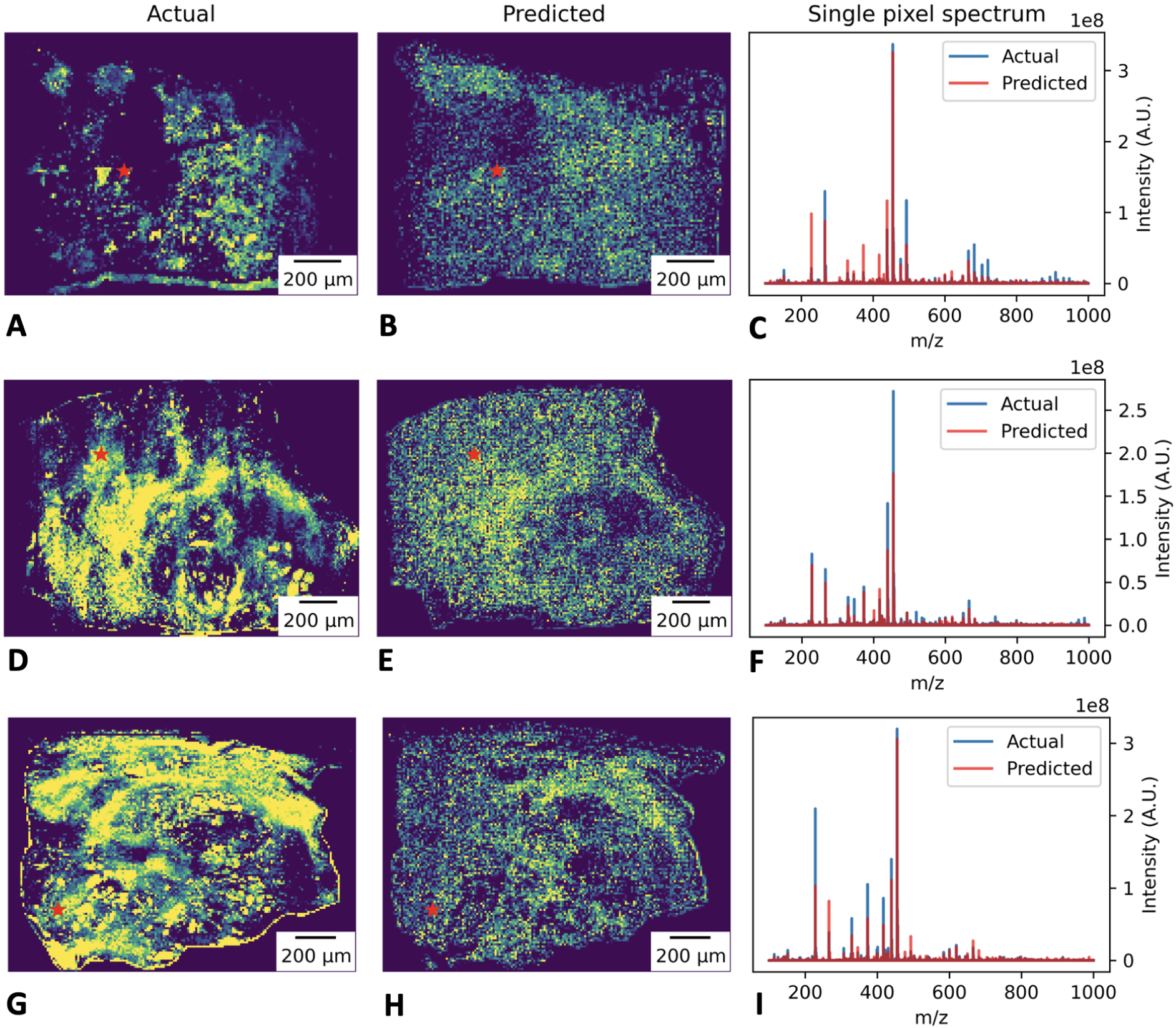
Prediction of MSI directly from regridded H&E features. **(A), (D)**, and **(G)** show the first PCA components of MSI for three different tissue specimens. **(B), (E)**, and **(H)** shows the corresponding predicted MSI PCA component directly from H&E. For the three tissue specimens, spectra of a single pixel (red star marker) from the cancerous region are shown both for actual and predicted images in **(C), (F)**, and **(I)**. Negative values of the predicted spectra are clipped here. Same range of intensity values in arbitrary units is used in this figure for each predicted image. Similar patterns can be seen in the predicted MSI images directly from H&E corresponding to the actual MSI images, with row 2 having slightly better prediction compared to the other two rows. The accompanying predicted spectrum for each row is clearly able to capture most of the m/z peaks in the actual spectrum, although there is an intensity mismatch between the actual and predicted spectrum.

From the *m/z* features, we obtained for MSI peak prediction as discussed in the methods section, we found that ground truth MSI image for the first tissue specimen with *m/z* 782.5655, which is identified as phosphatidylcholine PC(34:1) (Δ ppm = 1.93), reveals a cancerous region as shown in Figure 5B. The predicted MSI for this biomarker appears to identify cancerous regions in Figure 5C, when compared with the original tissue annotation in Figure 5A. We found another chemical biomarker *m/z* 780.5483, which is identified as cardiolipin CL(80:9) (Δ ppm = 0.18) that provided similar result as seen with the first tissue specimen. We tested and validated with another tissue specimen and achieved satisfactory result as the cancerous regions can be identified both by ground truth and predicted MSI images as shown in Figure 5D and Figure 5E respectively. Like the original MSI, the predicted MSI images for these two *m/z* values also show the same secondary predicted cancer regions that were not labelled by a pathologist. Figure 5 shows that in addition to labelling PC, our method provides useful additional information in the form of predicted chemical information. Since our cancer predictions are made directly from inferred chemical signatures, the results are inherently explainable by the chemical signatures at each point that were predicted from patterns detected on the H&E images.

**Figure 5:**
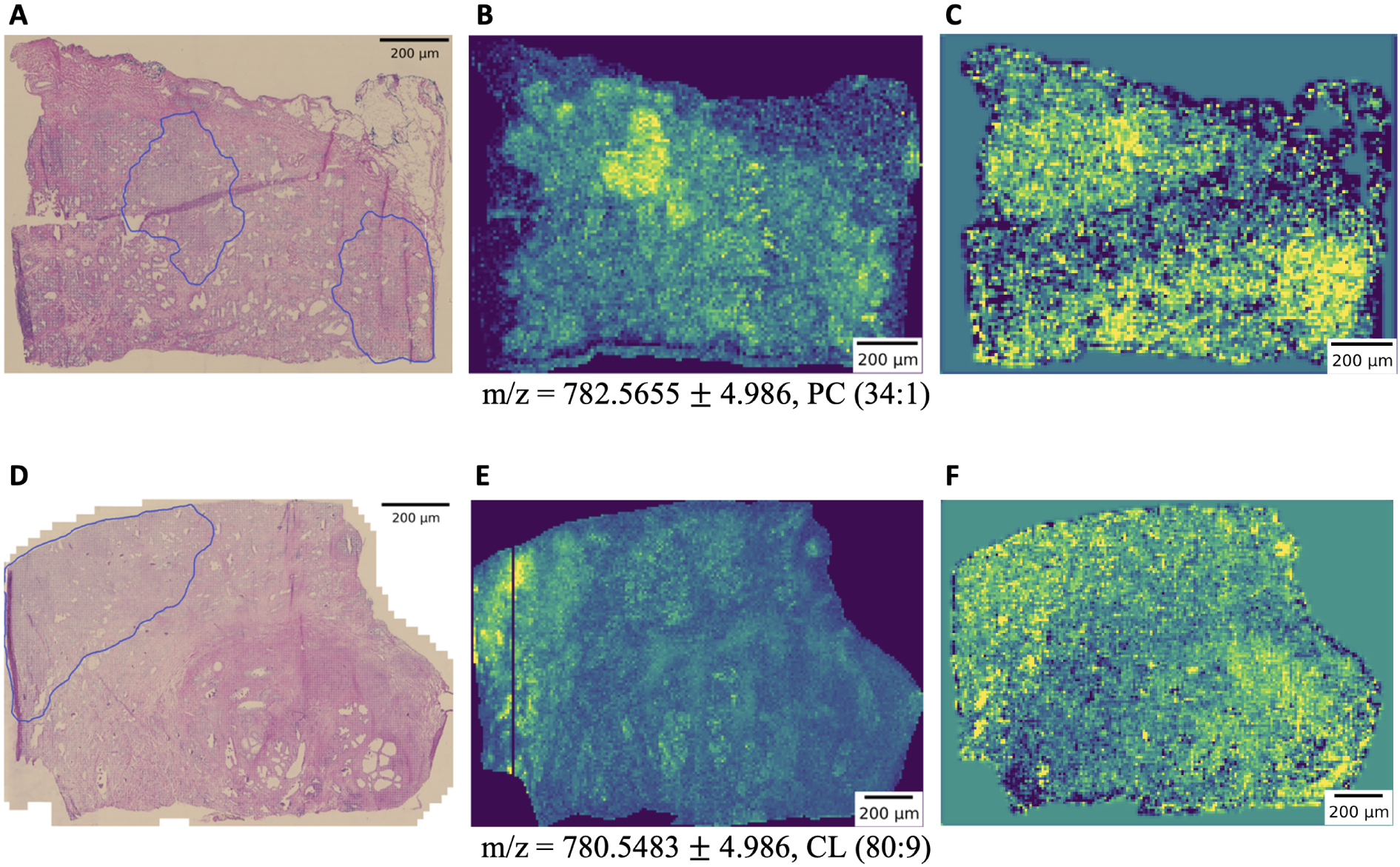
Comparison of cancer regions prediction directly from H&E with ground truth MSI. **(A)**, and **(D)** are the original cancer annotation for two different tissue specimens. **(B)** is the ground truth MSI of the first tissue specimen for *m/z* 782.5655 which is identified as phosphatidylcholine PC(34:1) (Δ ppm = 1.93). **(C)** is the predicted MSI of the first tissue specimen directly from H&E for the same *m/z* as ground truth. **(E)** is the ground truth MSI of the second tissue specimen for *m/z* 780.5483 which is identified as cardiolipin CL(80:9) (Δ ppm = 0.18). **(F)** is the predicted MSI of the second tissue specimen directly from H&E for the same *m/z* as ground truth.

## 4 Discussion

In this work, we have developed a machine learning approach to detect PC directly from H&E data incorporating chemical information found in MSI data. We found that H&E can predict mass spectra somewhat accurately, indicating a correlation between features visible in optical H&E imaging and the chemical information present in MSI. We also found that prostate cancer regions can be predicted reliably from MSI, outperforming H&E-based prediction, indicating that the mass spectra contain useful information for segmentation of cancerous regions. However, since MSI data are expensive to acquire and unavailable in a typical pathology lab, direct use of MSI data is infeasible in practice.

Instead, used paired MSI and WSI data for the five tissue samples to show a proof of principle that such paired data can be useful for training feature extraction networks on WSI data. We verified that the overlapping information between modalities matches that needed for segmentation, by predicting cancerous regions directly from H&E as well as from predicted MSI. Moreover, we found two MSI biomarkers (Figure 5) corresponding to specific masses that correctly identified the cancerous regions. Our prediction using H&E and MSI data was able to identify those regions as well.

As shown in the results section, secondary regions are also identified in some cases using MSI. These regions could be errors due to random noise or cancerous regions which were missing in the original pathology annotation. In future studies, additional validation of these secondary regions would be useful, for example using immunohistochemistry (IHC) imaging.

Our approach shows the feasibility of using readily available H&E data to predict the rich chemical information available in MSI images. Although our training process requires paired data including both H&E and MSI data from the same samples, the resulting trained models could be relevant in clinical settings where only H&E is available. To date, large public datasets of H&E images have been collected to support automated cancer detection and diagnosis, including manually curated slide-level or pixel-level annotations acquired at great cost. Our results suggest that we can reliably train feature extraction networks for automated WSI-based pathology without the need for time-consuming and expensive manual annotation by expert pathologists, as we achieve similar results with both MSI supervision and pathologist supervision. Even with our small sample set, we find that this approach gives reasonable results to show that PC could potentially be diagnosed accurately with correlating MSI and WSI. However, we do expect that accuracy of our predictions could be improved in future work by using deep learning with end-to-end training in a semi-supervised approach along with large-scale datasets such as Panda [21], which has around 11,000 whole-slide images of digitized H&E-stained biopsies, none of which is paired with MSI. These preliminary results along with a relative lack of MSI in public pathology datasets motivate the collection of larger paired H&E/MSI datasets in the future to support large-scale feature learning efforts for H&E analysis. Overall, we have demonstrated that we could potentially improve PC detection from H&E slides by incorporating MSI data, and that augmentation of labelled H&E datasets with paired MSI data can improve explainability by providing chemical information to support predictions. This lays the groundwork for developing more accurate prediction of mPC in patients to improve patient care.

## Acknowledgments

This research is supported by the Office of Research and Development, Veterans Health Administration, award MVP017. This publication does not represent the views of the Department of Veteran Affairs or the United States Government. This manuscript has been authored by UT-Battelle, LLC under Contract No. DE-AC05-00OR22725 with the U.S. Department of Energy. The authors also wish to acknowledge the support of the larger VA partnership, NIH grants U54-CA210180 (N.Y.R.A.), P41-EB028741 (N.Y.R.A.), and T32EB025823 (S.A.S.).

